# Rocking around the pheno-clock: bridging vegetation phenology and chronobiology

**DOI:** 10.64898/2026.02.24.707703

**Authors:** Bajocco Sofia, Ricotta Carlo, Bregaglio Simone

## Abstract

Plant phenology controls resource acquisition, survival and reproductive success, yet it is still commonly reduced to a limited set of calendar-based metrics, such as discrete dates of budburst or senescence. By contrast, chronobiology quantifies biological rhythms from full activity profiles using metrics that capture timing, regularity and amplitude of rest–activity cycles. Here, we bridge these two perspectives by developing a pheno-clock framework that translates plant phenology into chrono-ecological properties. We applied actigraphy-inspired metrics to multi-year satellite time series of European beech (*Fagus sylvatica*) forests across their European range. From daily photothermal activity profiles, we derived indices describing the strength, fragmentation and amplitude of annual rest–activity rhythms and related them to classical phenological metrics and regional climate. Our results reveal a marked asymmetry between spring and autumn phenology, indicating that canopy decline is governed by the cumulative organization of annual energy input, whereas canopy activation is dominated by short-term forcing. Across biogeographical regions, beech forests segregate into distinct pheno-chronotypes that differ in the timing and consolidation of rest and activity phases rather than in growing-season length alone. These chrono-ecological patterns suggest that climate filters not only when forests grow, but also how they structure their annual rhythmicity. By importing chronobiology into plant phenology, the pheno-clock framework provides a transferable approach to describe and compare seasonal strategies in plants, opening new avenues to link phenological diversity, functional traits and ecosystem responses to environmental change.

## 1. Introduction

Biological timing is a fundamental organizing principle across life, synchronizing internal physiology with external environmental cycles (Arendt, 1998; Hut et al., 2012). Time can be viewed as a functional dimension that shapes how species interact with and respond to their environment, defining their ecological temporal niche (Hut et al., 2012). The emerging “ecology of biological rhythms” explores how internal clocks and their rhythms, from circadian to circannual scales, mediate organism–environment interactions, influence behavioural strategies and competitive outcomes, and ultimately affect survival (Helm et al., 2013; Hut et al., 2012). When these rhythms become misaligned with environmental cues or with the timing of interacting species, phenological mismatches can arise with far-reaching consequences for fitness and ecosystem functioning (Renner & Zohner, 2018; Thoré et al., 2024; Terasaki Hart et al., 2025).

In humans, the circadian system coordinates daily cycles of physiology and behaviour, entrained to the 24-h day by external Zeitgebers (i.e., time-keepers) such as light, temperature and food intake (Roenneberg & Merrow, 2016; Quante et al., 2019). In deciduous trees, an analogous system operates at the annual scale: circannual rhythms organize transitions between dormancy, budburst, canopy development and senescence (Rohde & Bhalerao, 2007; Singh et al., 2017). These seasonal transitions are not simple passive responses, but reflect endogenous, anticipatory programmes that evolved to optimise resource allocation, stress avoidance and reproductive success in temperate climates. Their phase and amplitude are tuned by external Zeitgebers, primarily photoperiod and temperature, and their synchronisation with these cues is critical for individual performance and ecosystem functioning (Körner & Basler, 2010; Chuine et al., 2016; Fu et al., 2019).

Plant phenology thus encapsulates the temporal dimension of plant ecology and is highly sensitive to ongoing climate change (Piao et al., 2019). Yet most phenological studies still describe seasonality using a limited set of calendar-based events, such as budburst, leaf senescence or growing-season length, rather than as the continuous oscillation of a biological clock (Morellato et al., 2010). This event-based view obscures the recurrent, cyclic nature of plant time and limits our ability to compare rhythms among individuals, populations and regions. By contrast, chronobiology routinely characterizes internal rhythmicity from continuous rest–activity profiles, using standardized metrics of timing, regularity and amplitude (Refinetti, 2016; Roenneberg & Merrow, 2016). Actigraphy, which infers sleep–wake patterns and daily activity rhythms from continuous movement recordings, has become a key tool to quantify chronotypes and assess rhythm consolidation or fragmentation in humans (Gao et al., 2023).

Deciduous forests offer a natural parallel to the human rest–activity cycle. Over the annual cycle, trees alternate between phases of intense photothermal activity, dominated by photosynthesis, transpiration and carbon assimilation, and phases of metabolic quiescence during dormancy. This analogy suggests that tools developed for circadian rest–activity analysis could be adapted to quantify the timing, amplitude, and fragmentation of plant seasonal activity, and to define “pheno-chronotypes”, the temporal counterparts of human chronotypes, along early–late continua (Chuine et al., 2016). Such temporal diversity within species may represent an underexplored axis of phenotypic plasticity and adaptive potential under changing climatic conditions (Bonamour et al., 2019; Kramer et al., 2017).

The evolution of plant phenology research has been tightly linked to advances in observation. Satellite-derived vegetation indices such as NDVI and EVI (Rouse et al., 1974; Huete et al., 2002) have enabled large-scale monitoring of greening, dormancy and productivity (Zhang et al., 2003; Liu et al., 2015), but they typically compress complex physiological processes into smooth temporal curves. As a result, they provide only a coarse description of how plants distribute activity and rest across the year. Recently, process-based approaches have begun to bridge this gap. The SWELL model, for example, decomposes vegetation index time series into daily photothermal response curves, yielding a functional representation of plant activity throughout the annual cycle (Bajocco et al., 2025). This opens new opportunities to study circannual rhythmicity and its modulation by environmental drivers from space-borne observations.

Here, we build on these developments to propose a “pheno-clock” framework that translates deciduous forest phenology into a rest–activity language inspired by human chronobiology. We combine SWELL-derived photothermal activity with non-parametric actigraphic indices to describe the structure, stability and fragmentation of circannual rhythms. We apply this framework to long-term satellite time series of European beech (*Fagus sylvatica*) across major biogeographical regions to (i) characterise annual rest–activity patterns using chronobiology-based actigraphic indices; (ii) relate these chrono-ecological properties to classical phenological phases; and (iii) identify regional pheno-chronotypes, from “early risers” to “late sleepers”. By importing chronobiological concepts into plant phenology, we aim to provide a unified way to quantify how forests experience, structure and potentially adapt their temporal organization under a changing climate.

## 2. Data

European beech occurrence data were obtained from the high-resolution pan-European dataset by Strona et al. (2016), available at (Arendt, 1998) (Figure 1).

**Figure 1.**
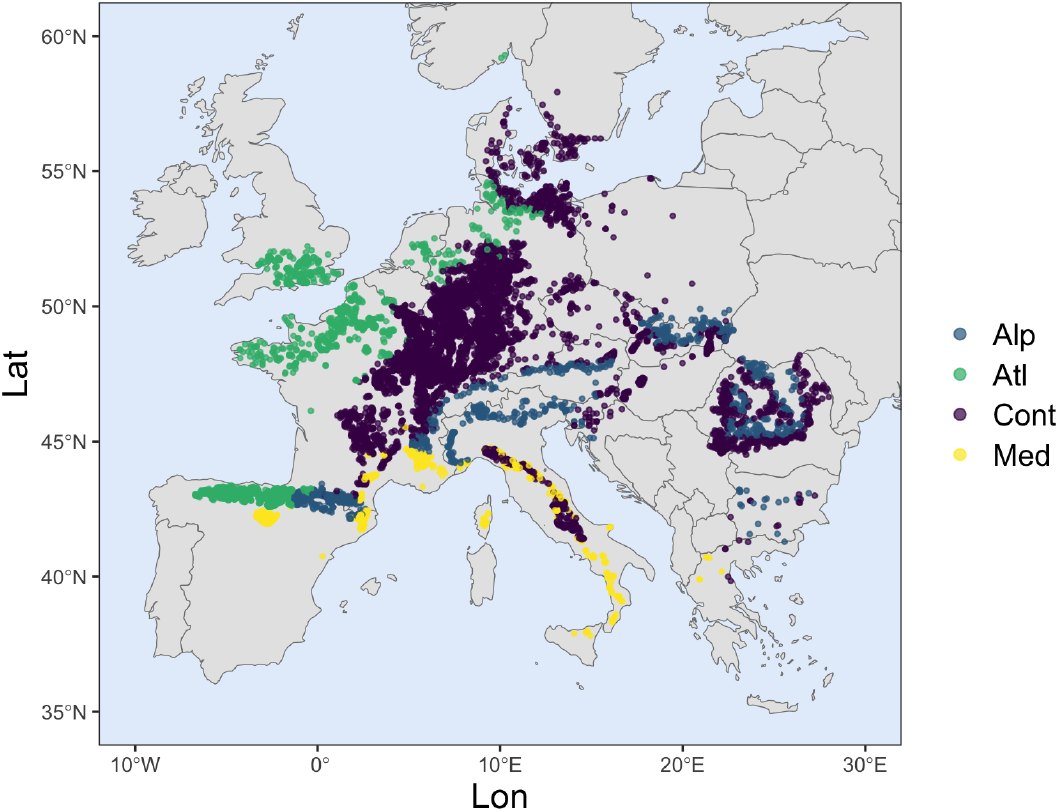
European beech forest locations across biogeographical regions: Alpine (Alp), Atlantic (Atl), Continental (Cont) and Mediterranean (Med).

Daily maximum and minimum air temperatures (0.1° resolution) for 2003–2023 were sourced from the E-OBS Copernicus dataset (https://www.ecad.eu), with hourly series estimated for chilling accumulation according to Campbell (1985).

Vegetation dynamics were derived from MODIS Terra and Aqua EVI (Enhanced Vegetation Index) products (250 m) via Google Earth Engine (Gorelick et al., 2017). Sixteen-day maximum value composites (Holben, 1986) were merged into 8-day profiles (46 per year) and smoothed using Savitzky-Golay filtering (Chen et al., 2004), with a window half-width of 5 and a sixth-degree polynomial. To ensure spatial consistency between MODIS EVI, E-OBS weather data, and beech occurrence points, we excluded single-tree, fragmented, and non-forest sites, retaining only points within the natural beech range (EUFORGEN; https://www.euforgen.org/species/fagus-sylvatica) and CORINE Land Cover 2018 (https://land.copernicus.eu/en/products/corine-land-cover/clc2018) deciduous forest class, resulting in 10,829 observations. Biogeographical regions were assigned using the map of the global biogeographical regions ("https://www.eea.europa.eu/en/datahub/datahubitem-view/11db8d14-f167-4cd5-9205-95638dfd9618). Four main biogeographical regions were considered for the analysis: Alpine (Alp), Atlantic (Atl), Continental (Cont) and Mediterranean (Med) (Figure 1).

## 3. Methods

### 3.1 Modelling annual photothermal activity

To derive daily-scale annual EVI profiles for beech forest locations, we applied the SWELL model (Simulated Waves of Energy, Light, and Life; Bajocco et al., 2025) to observed MODIS EVI time series. SWELL is a process-based phenology model that represents vegetation dynamics as a sequence of phenophases (dormancy, growth, maturity, senescence) governed by species-specific photothermal (PT) response functions. By fitting the observed EVI profiles with SWELL, we obtained smoothed daily trajectories that capture the timing and progression of key phenophases, as well as the daily course and magnitude of PT response rate, throughout the annual cycle (Figure 2). Considering the annual PT response derived from SWELL as the vegetation analogue of human daily physical activity measured by the actigraphy, we converted it into a “PT activity” signal suitable for rest–activity analysis. To mirror the progressive reduction of activity during senescence and the transition to rest during dormancy, the PT response was re-computed for the final stages of decline and for dormancy (Figure 2). During the growth, greendown, and initial decline (decline percentage < 49%) phases, the PT activity was directly assigned from the corresponding SWELL rate variables (growth rate, greendown rate, and decline rate, respectively). In contrast, for the final stage of decline (decline percentage ≥ 49%) and the dormancy induction phase, PT activity was defined as inversely related to the completion of the process, and calculated as 1 – decline rate and 1 – dormancy induction rate, respectively. The PT activity during the dormancy phase was determined using a two-step conditional logic. First, the temperature-dependent endodormancy response was computed: this response was set to zero when daily minimum temperatures were ≤ 3.4°C, and otherwise defined as 1 – endodormancy rate. Second, the endodormancy response was combined with the ecodormancy rate according to their co-occurrence: when both were active, their mean was used; when only one was active, its value was retained; and when neither was active, the PT activity was set to zero.

**Figure 2.**
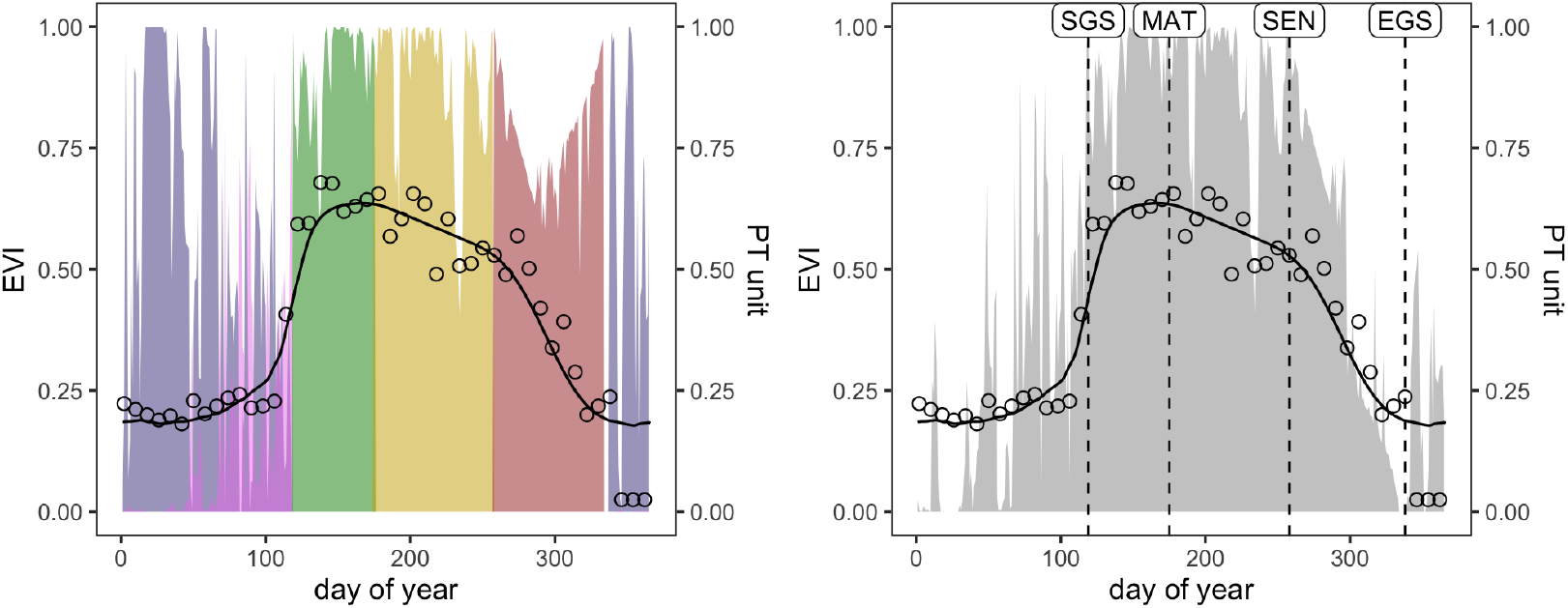
Example of the annual distribution of (left) daily photothermal (PT) response (coloured areas) and (right) its conversion into PT activity (grey area) across phenophases: endodormancy (purple), ecodormancy (violet), growth (green), greendown (gold), decline (red). Dashed vertical lines indicate the onset of the corresponding phenophases: start of growing season (SGS), maturity (MAT), senescence (SEN) and end of growing season (EGS). The solid black line indicates the EVI profile simulated by SWELL, while black dots indicate MODIS EVI observations.

### 2 Actigraphic analysis

The PT activity time series derived for the period 2003-2023 was used as input for the actigraphic analysis. To enable the application of standard chronobiological analytics, we rescaled each annual cycle (365 days) into one day (24 hours), so that 15 days approximately correspond to one hour, and one day to four minutes. The resulting hourly-scaled annual PT activity profiles were analysed using the R package ‘nparACT’ (Blume et al., 2016), which computes non-parametric actigraphic indices originally developed in chronobiology (Table 1). From these profiles, we derived two groups of indices adapted to characterize the annual cycle of beech forests. The first group comprises acti-metrics, including Relative Amplitude (RA), Inter-daily Stability (IS), Intra-daily Variability (IV), average activity during the ten consecutive hours of Maximum activity (M10) and five consecutive hours of Least activity (L5). The second group comprises acti-phases, defined as the start time of the M10 and L5 periods. Together, these indices describe the amplitude, stability, and continuity of the annual vegetation rest-activity cycle.

**Table 1.** Definition and meaning of the actigraphic indices and their translation from circadian to circannual rhythm.

|  | **Actigraphic indices** | **Original Definition (circadian rhythm)** | **New Definition (circannual rhythm)** |
| --- | --- | --- | --- |
| Acti-metrics | IS | Inter-daily stability | Inter-annual stability of photothermal (PT) activity |
|  | IV | Intra-daily variability | Intra-annual variability of PT activity |
|  | M10 | Mean activity during the most active 10-hour period | Mean PT activity during the most active 5-month period |
|  | L5 | Mean activity during the least active 5-hour period | Mean PT activity during the least active 2.5-month period |
|  | RA | Relative amplitude between L5 and M10 activity values | Relative amplitude between L5 and M10 PT activity values |
| Acti-phases | M10 start | Hour:minute corresponding to the start of the M10 period | Day of year corresponding to the start of the M10 period |
|  | L5 start | Hour:minute corresponding to the start of the L5 period | Day of year corresponding to the start of the L5 period |

RA is a non-parametric index that quantifies the contrast between periods of highest and lowest activity within a cycle (day, eq. 1).

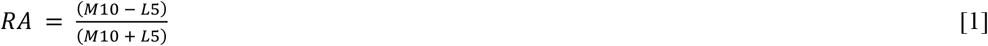

RA values were obtained by first calculating minute-level averages of activity across days. These minute-wise values were then used to compute rolling averages over all possible 10-hour (M10) and 5-hour (L5) windows within the 24-hour cycle (e.g., 6:00–16:00, 6:01–16:01, etc.). From these windows, the highest 10-hour mean (M10; i.e., the activity peak) and the lowest 5-hour average (L5; i.e., the minimum activity during rest) were identified together with their respective start times. When applied to the circannual cycle of PT activity in deciduous vegetation, the RA measures the contrast between maximum seasonal activity (equivalent to M10, the most active period) and minimum seasonal activity (by default equivalent to L5, the least active period), thereby reflecting the strength of annual amplitude. High RA values indicate strong seasonal amplitude and a clear separation between active growth and rest phases. The start times of M10 and L5 were initially expressed into hour:minute format and subsequently converted to day-of-year (doy) values before subsequent analysis. To avoid distorting statistical analyses and to ensure temporal continuity for acti-phases crossing the calendar year, L5 doy values occurring after 31 December were recoded by adding 365 (e.g. 1 January = doy 366, 2 January = doy 367, and so on) up to doy 100. IV captures the degree of fragmentation within a rest-activity rhythm and was computed according to eq. 2.

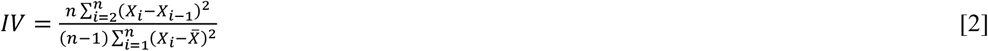

where n is the total number of sampling points, 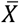 is the grand mean of the time series and X_?_ denotes the activity value at each sampling point (hourly values). IV approaches zero for a smooth sinusoidal signal and increase towards two under Gaussian noise. When applied to vegetation, IV captures the degree of fragmentation of seasonal dynamics, and reflects irregularities or fluctuations in the seasonal curve, such as interruptions caused by late frost, heatwaves, or other disturbances. A smooth growth–senescence pattern is therefore associated with low IV, whereas high IV indicates noisy or multi-modal seasonal signals.

IS measures the stability of rest-activity rhythms and reflects how consistently a given pattern is repeated across cycles (eq. 3).

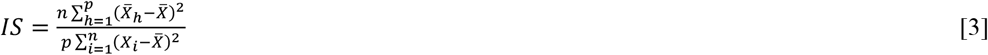

where p is the number of sampling points per day, and 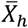 represents the mean activity at each sampling time (hourly means). IS ranges from 0, corresponding to Gaussian noise, to 1, with higher values indicating stronger rhythmic stability and entrainment to an annual Zeitgeber. In the vegetation context, IS quantifies the consistency of PT activity rhythms across years and reflects how reliably external environmental drivers synchronize beech PT activity and phenology to annual cycles. Values closer to 1 indicate stronger alignment with environmental cues such as photoperiod and temperature.

To explore differences in PT activity patterns among biogeographical regions, we carried out a Principal Component Analysis (PCA), using acti-metrics (IV, IS, RA, M10 and L5) and acti-phases (L5 and M10 start times) as input variables. We then conducted region-specific Spearman rank correlation analyses between actigraphic indices and SWELL-derived phenophases to quantify the relationships between beech forests phenology and annual photothermal rhythms. To assess whether the timing of maximum (M10) and minimum (L5) PT activity differed significantly from classical phenological phases, we performed paired t-tests within each region comparing M10 start time with the start of the growing season (SGS), and L5 start time with the end of the growing season (EGS). Finally, to identify the “pheno-chronotype” of the different biogeographical regions, we translated the acti-phases into a “pheno-clock” and carried out a paired t-test between all combinations of biogeographical regions, separately for the start times of the highest (M10) and lowest (L5) PT activity periods.

## 4. Results

### 4.1 Rest-activity biogeographical patterns

The Principal Component Analysis (PCA) identified the acti-graphic indices that best characterize European beech forests across biogeographical regions, with the first two components capturing 74.1% of the total variance (PC1 = 53.7%, PC2 = 20.4%; Figure 3). PC1 mainly represented a gradient from high-amplitude, stable rhythms with deep rest (low L5, high RA, high IS, low IV) to low-amplitude, fragmented rhythms with elevated baseline activity (high L5, low RA, low IS, high IV). PC2 primarily contrasted sites according to the magnitude and timing of maximum and minimum activity, opposing M10 values to the timing of both M10 and L5 onsets.

**Figure 3.**
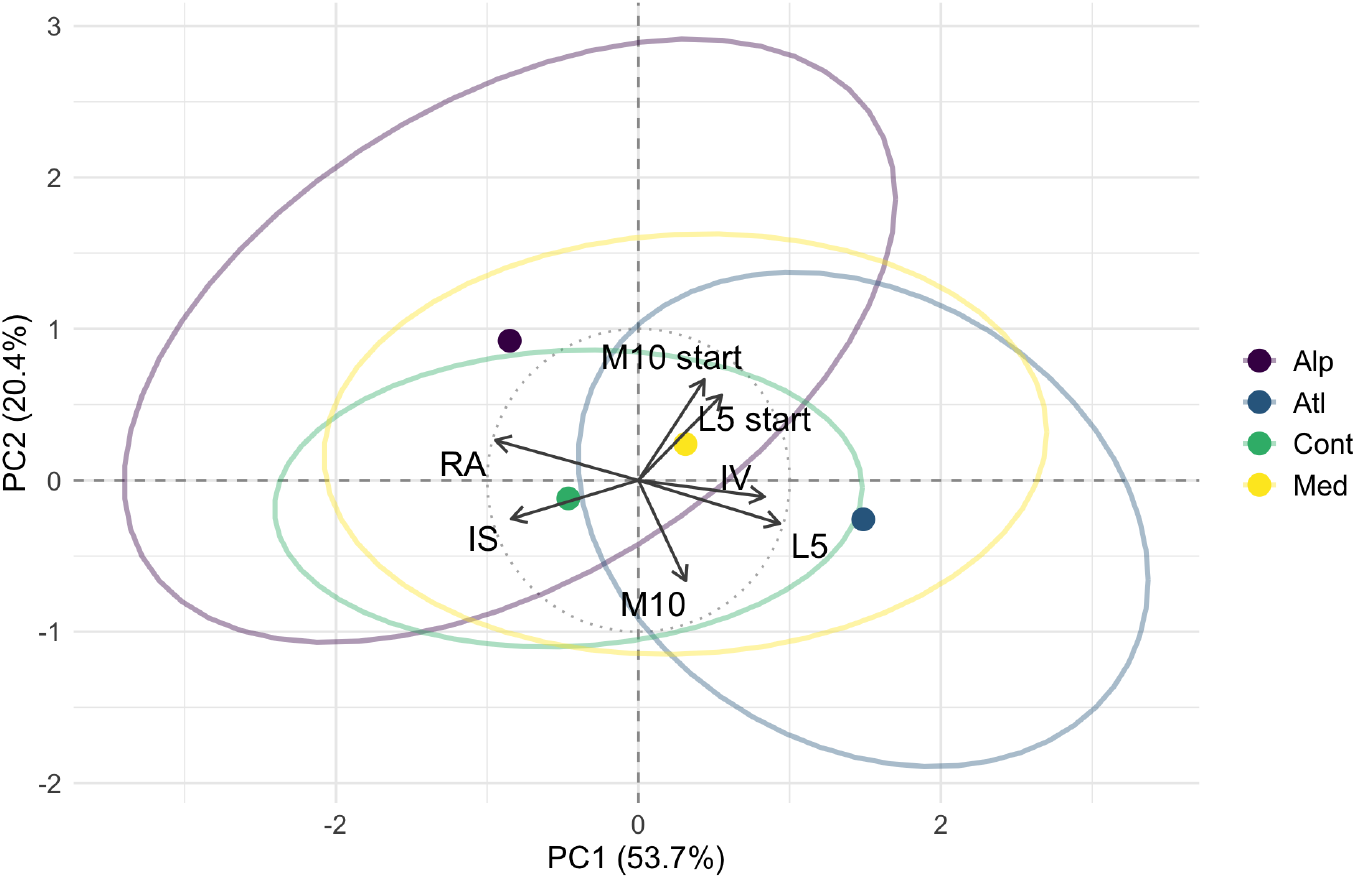
Principal Component Analysis (PCA) biplot of actigraphic indices across biogeographical regions. Regions are represented by their centroids and 50% confidence ellipses. Arrows represent variable loadings: RA (relative amplitude between L5 and M10); IS (inter-annual stability); IV (intra-annual variability); M10 (mean photothermal activity during the highest 10-hour period); L5 (mean photothermal activity during the lowest 5-hour period); M10 start and L5 start (day of year of the corresponding start times).

Biogeographic regions’ centroids occupied distinct quadrants of the biplot. Atlantic sites were associated with high L5 and IV and low RA, indicating flatter, less contrasted and more fragmented annual cycles. Alpine sites showed the opposite pattern, with high RA and very low L5 and M10 values, reflecting strong seasonal contrast and sharply defined rest–activity phases. Mediterranean sites were characterized by a late onset of both L5 and M10 and low IS, pointing to delayed and less coherent rhythms shaped by extreme events and irregular recovery. Continental sites combined high IS with early onsets of L5 and M10, consistent with strong inter-annual coherence of the photothermal signal and a short, temperature-constrained growing season.

### 4.2 Correlating actigraphic indices and phenophases

Correlations between phenological phases and actigraphic indices were broadly consistent across regions although their strength varied (Figure 4). Senescence (SEN) and the end of the growing season (EGS) showed the tightest associations with the annual PT rhythm, being positively correlated with both maximum and minimum activity (M10, L5; up to ρ = 0.68 for SEN and ρ = 0.79 for EGS) and negatively correlated with relative amplitude (RA; down to ρ = –0.79). Accordingly, smoother, low-amplitude PT cycles tended to coincide with delayed SEN and EGS, whereas sharper, high-contrast rhythms were associated with earlier canopy decline. In contrast, maturity (MAT) showed weak and inconsistent correlations with actigraphic indices (|ρ| < 0.36), while the start of the growing season (SGS) displayed slightly stronger relationships. Later SGS was linked to higher RA and lower L5, especially in Alpine region (|ρ| up to 0.45), indicating a tighter coupling between spring phenology and PT dynamics in energy-limited climates and a weaker link in the Mediterranean region. Intra-annual variability (IV) and inter-annual stability (IS) further separated early-from late-season phases. Later SGS occurred where PT forcing was more stable and less fragmented (positive correlations with IS and negative correlations with IV). In contrast, late-season phases showed the opposite pattern: SEN and EGS correlated positively with IV and negatively with IS, suggesting that vegetation activity is prolonged under intermittently favourable but less predictable PT conditions, and compressed into a narrower seasonal window where climates are more stable. Correlations between acti-phases and phenophases were generally positive, indicating shared drivers of PT timing and phenology. The strongest relationships were between L5 start and EGS (up to ρ = 0.76 in Alpine and Continental regions), highlighting a tight coupling between PT decline and growing season termination in cold, seasonal climates. By contrast, M10 start was more closely related to SEN than to SGS, supporting a strong carry-over of PT activity into the late season, while SGS remained only weakly linked to acti-phase timing, consistent with its regulation by short-term thermal forcing and chilling requirements rather than by the broader annual PT cycle.

**Figure 4.**
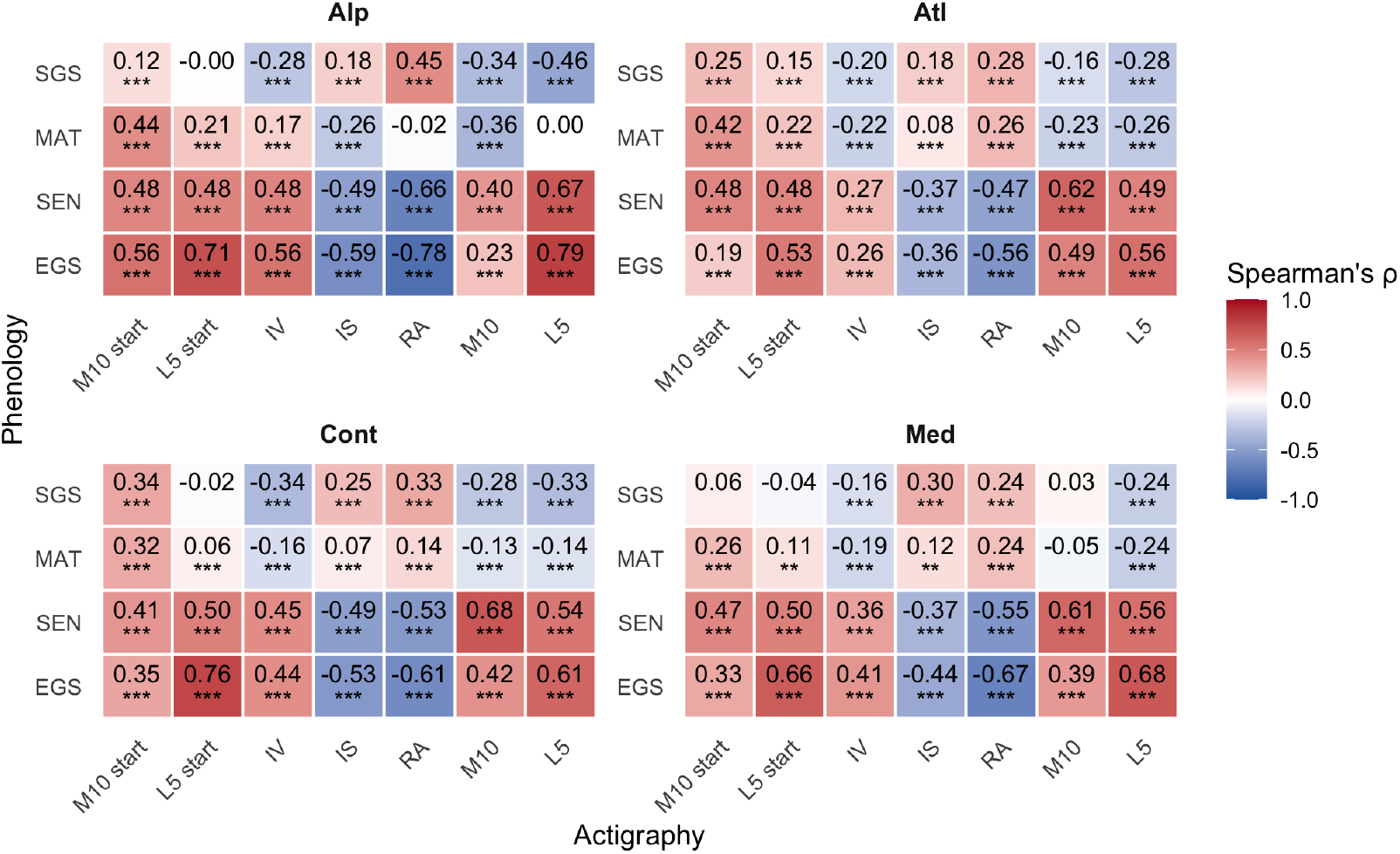
Spearman’s rank correlations (ρ) between actigraphic indices, (M10 start, L5 start, IV, IS, RA, M10 and L5) and phenological phases derived from SWELL (SGS, MAT, SEN, EGS) across biogeographical regions. Significance levels: * p < 0.05; **p < 0.01; *** p < 0.001.

### 4.3 Comparing photothermal activity and phenology

Annual photothermal activity showed a clear climatic signature, with marked regional differences in both amplitude and timing (Figure 5). Atlantic and Mediterranean regions displayed broader PT activity curves, indicating longer periods of favourable activity, whereas Alpine and Continental regions showed narrower curves, consistent with shorter, more concentrated growing seasons. Variability among sites was highest in Alpine and Mediterranean regions, suggesting stronger local constraints imposed by cold and drought, respectively. Across all regions, spring trajectories were more irregular, with frequent short-term fluctuations, while autumn decline was smoother, indicating stronger external forcing during growth than during senescence.

**Figure 5.**
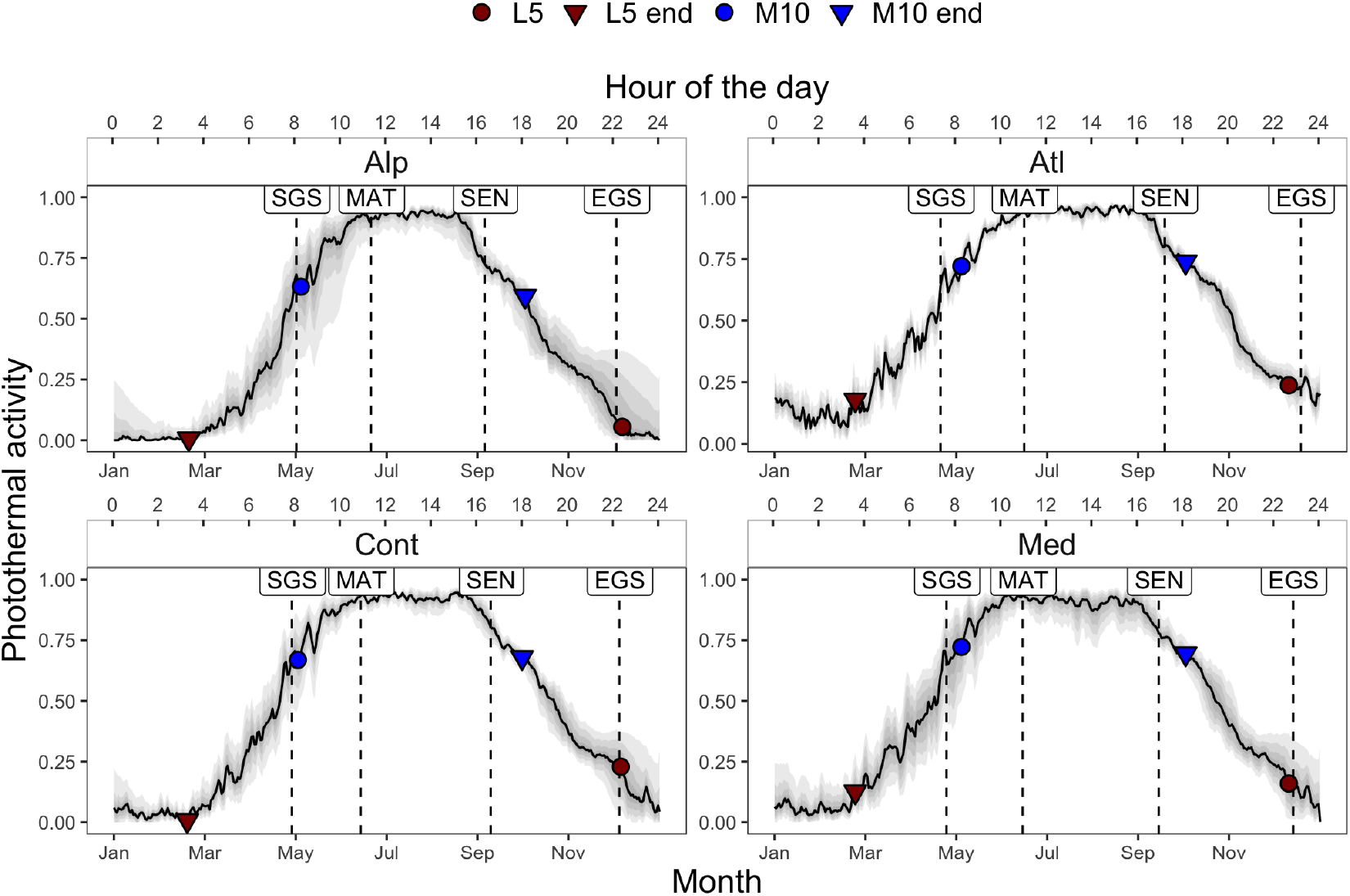
Annual mean profiles of photothermal activity in European beech forests across biogeographical regions, with shaded areas indicating inter-site variability. The mean start and end of the highest 10-hour (M10) and lowest 5-hour (L5) activity periods are shown as circles and triangles, respectively, coloured in blue (M10) and red (L5). Vertical dashed lines indicate the regional mean of the phenophases: start of growing season (SGS), maturity (MAT), senescence (SEN), and end of growing season (EGS).

The temporal alignment between actigraphic and phenological phases further highlighted this pattern (Figure 5; Table 2). In Atlantic and Mediterranean regions, the onset of maximum activity (M10) started up to nearly three weeks after SGS, implying that conditions for intense PT activity are reached well after canopy green-up. This gap narrowed in Alpine and Continental regions, where SGS and M10 onset nearly coincided, indicating a shared, strong climatic control on spring dynamics. For minimum activity (L5), the mismatch between L5 onset and EGS was smaller across all regions and followed a clear gradient: L5 started about one week before EGS in Atlantic and Continental regions, roughly at the same time in Mediterranean sites, and about one week after EGS in Alpine sites. Paired t-tests confirmed that differences between acti-phases and phenophases were significant in all regions (p < 0.01; Figure 6).

**Figure 6.**
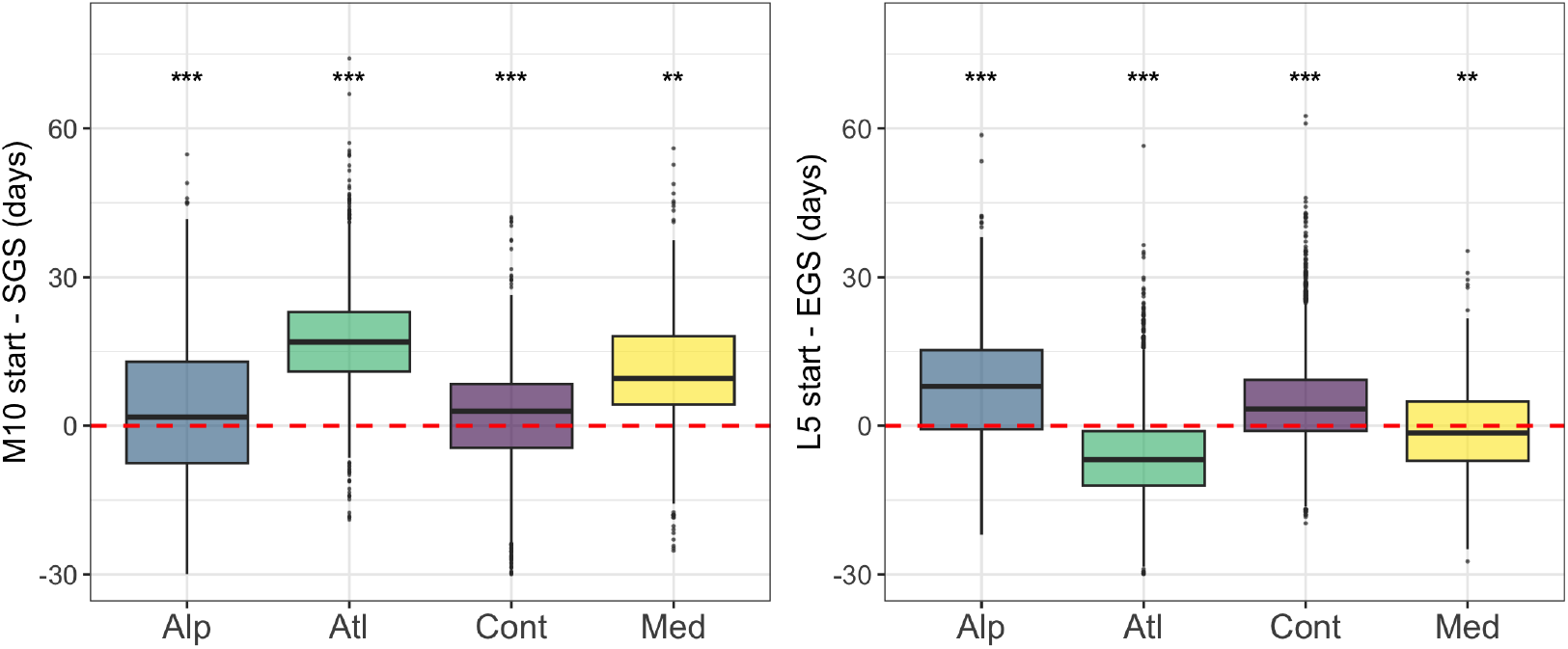
Boxplots showing the mean difference (in days) between actigraphic phases and the corresponding phenophases (SGS for M10; EGS for L5) across biogeographical regions. Positive values indicate acti-phases occurring after the phenophases, whereas negative values indicate earlier occurrence. The central line within each box represents the median difference; the box spans the interquartile range (IQR, 25th - 75th percentiles), and whiskers extend to 1.5 times × IQR. Values beyond this range are plotted as outliers. The red dashed line marks synchrony (0 days difference). Significance levels: * p < 0.05; **p < 0.01; *** p < 0.001.

### 4.4 Pheno-clock and chronotypes

The European beech forests “pheno-clock”, derived from the onset of the acti-phases, exhibited distinct patterns across the four biogeographical regions (Figure 7), revealing systematic shifts in the timing of PT activity along climatic gradients. The onset of the highest 10-hour activity period (M10 start) had a global mean start time of approximately 8:15 AM (corresponding to May 2-3), with a lag of approximately 25 minutes (roughly 6 days) between the earliest and the latest region. A clear biogeographical trend emerged: the Continental region, representing the “early riser” pheno-chronotype, began this active phase earliest (approx. 8:10 AM; May 1-2). In contrast, the Alpine, Mediterranean, and Atlantic regions exhibited progressively later onsets (approx. 8:25 AM to 8:30 AM; May 5-6 to May 7-8), with Atlantic beech forests representing the “late riser” pheno-chronotype. The onset of L5, which indicates the beginning of the prolonged and maintained “sleep” phase, showed much smaller regional differences. L5 timing varied by only about 10 minutes (equivalent to 2-3 days) across regions, centered on a global mean of approximately 10:45 PM (December 6-7). Also in this case, the timing followed a thermal gradient, with earlier onset in the colder Continental region and later onset in the warmer Mediterranean region, with Alpine and Atlantic in between.

**Figure 7.**
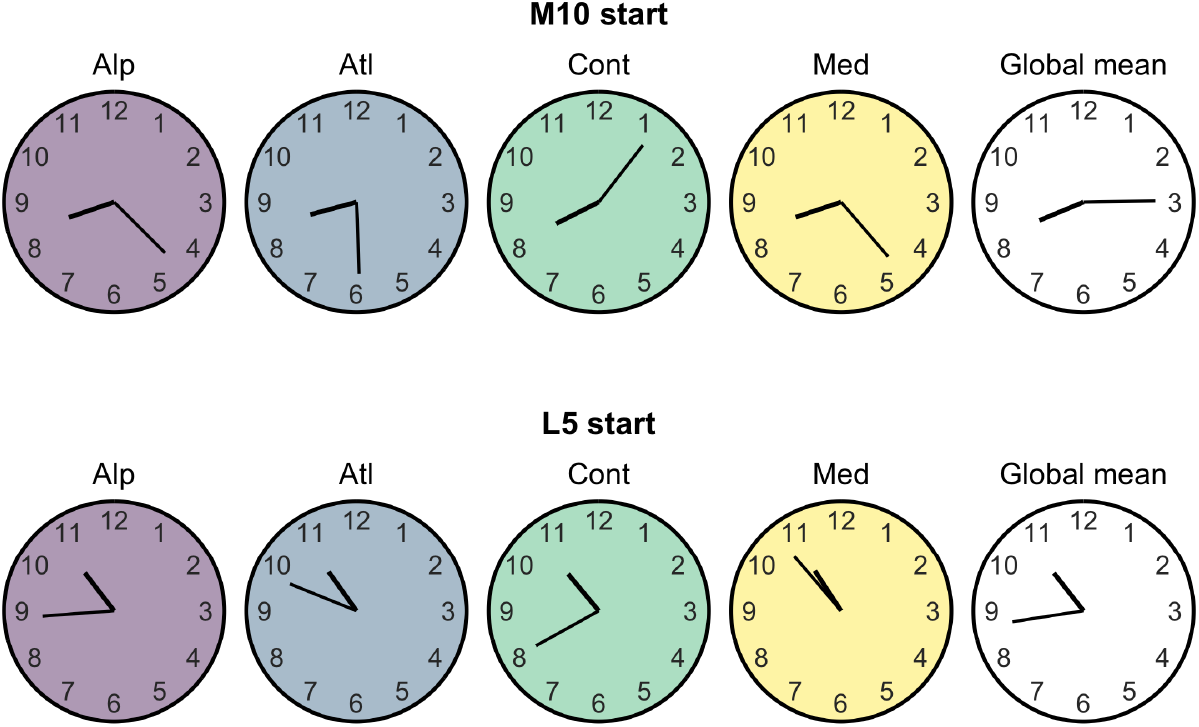
Pheno-clocks showing the start time of actigraphic phases using a 12-hour clock representation. The short hand indicates hours and the long hand indicates minutes. Start time for the highest 10-hour activity (M10) is AM, whereas start time for the 5-hour lowest activity (L5) is PM.

These pheno-clock patterns were statistically supported by the paired t-test comparing acti-phases between biogeographical regions (Figure 8). All regional differences were highly significant, except for M10 start between Alpine and Mediterranean and for L5 start between Alpine and Atlantic (and Mediterranean) regions. This result indicates that, although some beech forests are synchronized in the timing of maximum or minimum photothermal activity, they remain desynchronized with respect to the corresponding phenological phases, highlighting a decoupling between activity timing and phenophase occurrence.

**Figure 8.**
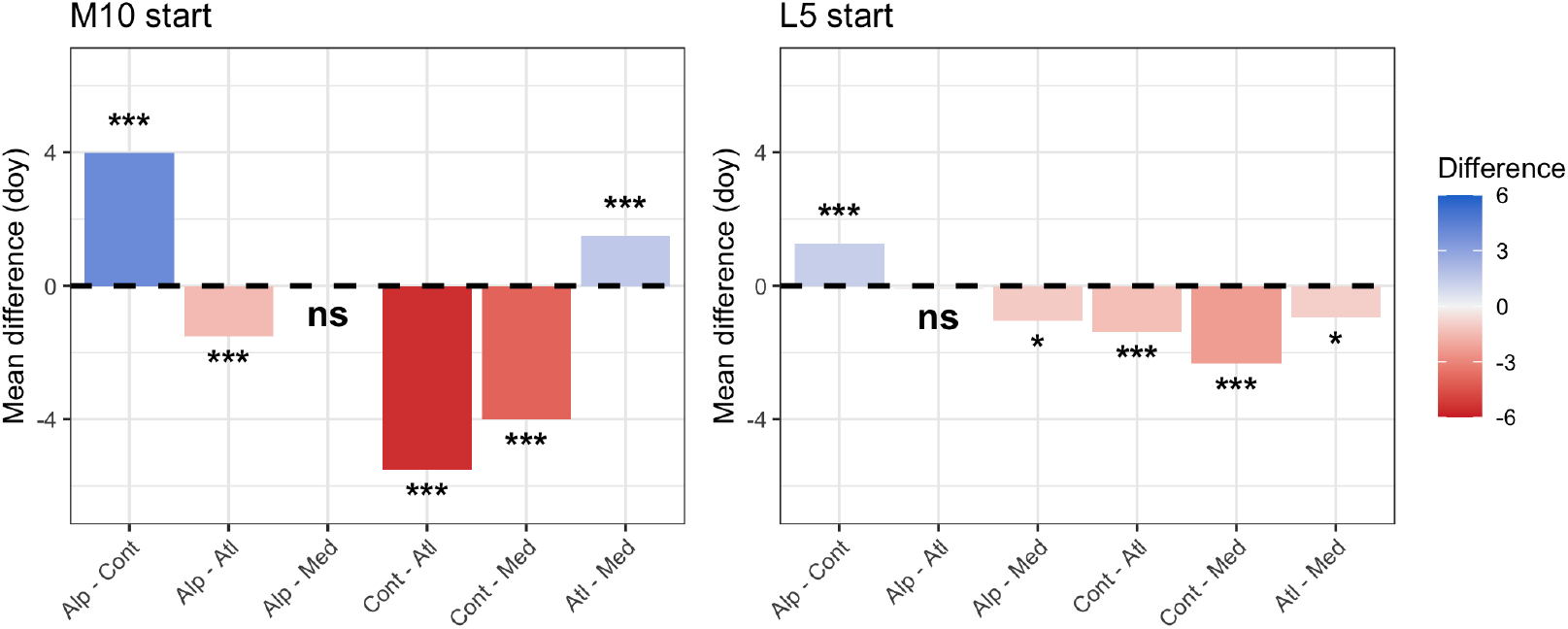
Histograms comparing the mean differences in actigraphic phase timing (day of year, doy) for M10 start and L5 start among biogeographical regions. Significance levels: * p < 0.05; **p < 0.01; *** p < 0.001; NS not significant.

## 5. Discussion

By reframing annual photothermal dynamics as a chronobiological rhythm, we show that European beech forests express contrasting activity–rest strategies across European climate gradients. Rather than simply shifting the timing of spring onset and autumn senescence, forests reorganize the internal structure of their annual cycles: how long activity or rest phases are consolidated, how abrupt transitions are, and how fragmented the rhythm becomes. Our actigraphy-inspired pheno-clock reveals that (i) rest–activity rhythms exhibit a clear biogeographical structure, with more consolidated cycles in cooler and more continental climates; (ii) photothermal rhythmicity is asymmetrically coupled to spring and autumn phenology; and (iii) beech forests segregate into distinct pheno-chronotypes that differ in temporal strategy rather than in growing-season length alone. These chrono-ecological properties add an explicitly rhythmic dimension to plant phenology and complement traditional event-based metrics (Piao et al., 2019; Delpierre et al., 2024).

### 5.1 Rest-activity biogeographical patterns

Actigraphic indices revealed a clear north–south and continental–oceanic gradient in the photothermal rhythm of European beech forests. Alpine and Continental regions showed high inter-annual stability, low intra-annual fragmentation and broad annual amplitude, indicating an early, sharp and well-consolidated transition between rest and activity under strong seasonal constraints. In contrast, Atlantic and Mediterranean regions exhibited flatter and less stable annual rhythms, with smoother and delayed transitions, consistent with more climatically buffered or transient conditions. These patterns match evidence that strongly seasonal climates support more predictable vegetation dynamics, whereas sites with lower seasonal coherence show greater irregularity (Rodriguez-Galiano et al., 2016; Škvareninová et al., 2024) and align with global NDVI-based analyses showing maximum amplitude and stability at high northern latitudes and minima in southern and arid regions (Recuero et al., 2019).

The combination of high IS and low IV in Alpine and Continental regions likely favours a tight coupling between climatic cues and phenological timing (Vitasse & Basler, 2012). Conversely, lower RA and higher L5 in Atlantic and Mediterranean regions point to weaker photothermal contrast and less consolidated rest. In Atlantic forests, this reflects oceanic climatic moderation that dampens seasonal thermal gradients (Karnieli et al., 2019; Terasaki Hart et al., 2025). In Mediterranean forests, high IV and low IS indicate fragmentation of the annual photothermal signal due to recurrent heat and drought stress, which can reduce productivity (Matula et al., 2023) and disrupt resource use and carbon assimilation (Gouveia et al., 2017; Ermitão et al., 2021; Zhang et al., 2024). Amplitude-related indices further suggest that Alpine and Continental beech forests maintain a broad, sharply defined seasonal rhythm with strong dormancy (Zani et al., 2020; Bajocco et al., 2025), whereas Atlantic and Mediterranean stands sustain higher baseline energy levels throughout the year, but at the cost of reduced seasonal contrast and potentially weaker dormancy induction (Vitasse & Basler, 2013; Baumgarten et al., 2021; Didion-Gency et al., 2024).

Overall, these results indicate that climate shapes not only the length of the growing season, but also the internal architecture of annual rhythms – namely, how consolidated or fragmented rest and activity are over time. Actigraphic indices therefore emerge as integrative temporal properties that respond to the same climatic drivers governing more conventional phenological and growth indicators (Vitasse et al., 2010; Beil et al., 2021; Yang et al., 2020).

### 5.2 Asymmetric coupling of photothermal rhythm and phenology

Across regions, our results reveal a clear asymmetry in the coupling between the photothermal (PT) rhythm and phenology in European beech forests. Correlations between PT timing and phenological phases strengthen from spring onset (SGS) towards the end of the growing season (EGS), indicating that the annual PT cycle is more tightly shaped by canopy decline than by spring activation.

This asymmetry is consistent with the mechanistic differences between spring and autumn phenology. SGS acts as a predominantly event-driven activation phase: budburst and early leaf expansion occur when short-term temperature forcing coincides with sufficient chilling and a permissive photoperiod (Wang et al., 2020; Walde et al., 2022; Jiang et al., 2024). In contrast, EGS emerges as a cumulative “decision point”. Senescence and the build-up of dormancy result from the integration of multiple cues operating over multiple seasons: progressive cooling, shortening photoperiod (Lang et al., 2024), leaf life-span constraints (Meier et al., 2025), legacy of spring–summer productivity (Keenan & Richardson, 2015; Rodriguez-Galiano et al., 2016; Zohner et al., 2023) and stress exposure (Mariën et al., 2021), closely tracking the seasonal decay of PT activity. The observed frequent short-term PT fluctuations in spring, versus the smoother autumn decay, further support this interpretation of spring as an event-driven, climatically unstable phase and autumn as a more cumulative, internally regulated one (Zhang et al., 2023; Zohner et al., 2023).

Under ongoing climate change, increasing weather instability is therefore likely to disrupt spring phenology more readily than autumn dynamics. Spring represents the most plastic and least synchronized phase of the annual cycle, whereas autumn remains more strongly constrained by photoperiod control and internal regulation. Yet, autumn canopy dynamics are particularly sensitive to climate variability and carry-over effects from prior conditions (Piao et al., 2019; Zani et al., 2020; Zhang et al., 2023; Zohner et al., 2023). From a chrono-ecological perspective, this implies that shifts in the timing, amplitude or consolidation of dormancy may alter the annual rhythm more profoundly than equivalent shifts in spring onset. Because autumn determines the transition into metabolic rest and the resetting of chilling and forcing requirements (Vitasse et al., 2014; Vitasse & Basler, 2013), changes in the rest phase can propagate forward, modifying the sensitivity of subsequent spring events (Fu et al., 2019; Baumgarten et al., 2021). The relatively weak imprint of SGS on annual rhythmicity should therefore not be interpreted as low ecological importance; rather it reflects the event-like nature of spring phenology, which is better captured by event-based metrics, whereas autumn and winter more strongly define the background rhythm against which such events occur.

### 5.3 Pheno-chronotypes and intra-specific temporal diversity

The timing of photothermal activity in European beech, viewed as a 24-h analogue of the annual growth–dormancy cycle, revealed distinct regional pheno-chronotypes. The onset of high (M10) and low (L5) activity phases point to contrasting temporal strategies for resource acquisition and stress avoidance. In Continental (and similarly Alpine) climates, both M10 and L5 start early, consistent with adaptation to a short, thermally constrained growing window. Here, beech forests express an “early chronotype”: they activate rapidly, reach peak activity early and enter “sleep” sooner, maximising carbon gain within a condensed yet predictable season while reducing exposure to early autumn frosts (Tylewicz et al., 2018; Charrier et al., 2021). Atlantic and Mediterranean forests instead behave as “late risers/late sleepers”, with delayed onsets of both M10 L5 start. This likely reflects the need to wait for stable weather conditions that allow sustained high PT activity, combined with prolonged favourable seasons that reduce selective pressure for early peak activity (Körner & Basler, 2010). Accordingly, weaker thermal constraints permit extended activity into autumn (Wingate et al., 2015), in line with the “late to bed, late to rise” dynamics suggested by Beil et al. (2021), where warm autumns delay senescence and dormancy and can lead to delayed leaf-out in the following spring (Fu et al., 2019).

The existence of regional pheno-chronotypes suggests that temporal diversity within species may contribute to resilience. Temporal strategies likely reflect both local genetic adaptation and phenotypic plasticity in response to intra- and inter-annual variability (Vitasse et al., 2010; Fu et al., 2019; Walde et al., 2022). Just as variation in human chronotypes provides population-level flexibility in coping with environmental and social rhythms (Roenneberg & Merrow, 2016; Helm et al., 2017), intra-species variation in PT activity timing may plastically buffer forest populations against climatic extremes (Bonamour et al., 2019). This evidence adds a temporal dimension to biodiversity that complements its spatial and functional components, suggesting that the diversity of rhythmic behavior itself may be an overlooked aspect of ecosystem resilience.

### 5.4 Ecological implications of the pheno-clock

Framing seasonal growth and dormancy as an annual rest-activity rhythm enables the development of metrics that quantify and compare plant behavior in new ways. In human chronobiology, actigraphic analyses are routinely used to assess rhythm amplitude, phase stability, and fragmentation, which serve as indicators of health and resilience (Smagula et al., 2015). When applied to deciduous forest dynamics, similar metrics could offer novel insights into vegetation ecological fitness. Amplitude-related metrics reflect the contrast between peak productivity and dormancy, while rhythm stability metrics reveal the predictability or plasticity of timing across years, key properties for efficient resource allocation and anticipation of environmental changes. These characteristics are especially relevant under climate change, where increased inter-annual variability and phenological mismatches may signal stress or maladaptation (Piao et al., 2019; Liu et al., 2018; Liu et al., 2025). For example, individuals or populations maintaining high rhythmic stability despite climatic fluctuations may exhibit greater resilience, whereas fragmented or erratic activity patterns may indicate higher vulnerability to stress (Clark et al., 2014; Montgomery et al., 2020; Jiang et al., 2025). Therefore, actigraphic metrics allow us to quantify not only when plants grow, but how robustly and consistently they engage in activity and rest over time, creating new opportunities to study phenotypic plasticity and local adaptation (Vitasse et al., 2010; Kramer et al., 2017; Bonamour et al., 2019).

Integrating plant phenology into a rest-activity architecture further supports the application of Zeitgeber theory, a cornerstone of chronobiology. Photoperiod, temperature forcing and chilling act as key Zeitgebers in forests (Basler & Körner, 2012; Renner & Zohner, 2018), and their regional variation and ongoing shifts under climate change directly influence the phase and amplitude of phenological rhythms. By conceptualizing phenology through the lens of entrainment and desynchronization, we gain a framework to analyze how internal clocks may fall out of step with the environment, leading to phase shifts (e.g., earlier budburst), amplitude suppression (e.g., reduced growing season), or altered phase relationships with internal (i.e., phenophases) or external dynamics (e.g., pollinators) (Forrest & Miller-Rushing, 2010).

The rest-activity perspective also enables regional comparisons of phenological dynamics that extend beyond average dates. While traditional approaches focus on temporal shifts in specific events (e.g., earlier budburst in southern regions), a rest-activity rhythmic framework captures whole-cycle properties: the duration and consolidation of activity and dormancy phases, the abruptness or smoothness of transitions, and the degree of fragmentation within the annual cycle. This supports the identification of new typologies of phenological strategies across climates and environments, and may help detect ecotonal shifts or phenological regime changes under warming scenarios.

From an analytical standpoint, translating seasonal plant dynamics into a rest-activity language permits the application of quantitative tools that are well-established in chronobiology but underutilized in plant science (Roenneberg & Merrow, 2016; Dominoni et al., 2023). This perspective shift moves phenology beyond discrete, timepoint-based analyses to a continuous and dynamic representation of plant activity, enabling a more holistic understanding of seasonal performance.

By embedding plant phenology within the ecology of biological rhythms (Hut et al., 2012; Roenneberg & Merrow, 2016; Refinetti, 2012; Thoré et al., 2024), the pheno-clock framework provides a quantitative and transferable approach to describe how forests organize their time. In a rapidly changing climate, understanding not only when forests grow and rest, but also how they structure their annual rhythms, is essential. The pheno-clock framework provides a first step in that direction, offering quantitative, transferable metrics to describe forest temporal strategies and their sensitivity to environmental change.

